# Opsin gene duplication in Lepidoptera: retrotransposition, sex linkage, and gene expression

**DOI:** 10.1101/2023.08.16.552946

**Authors:** Peter O. Mulhair, Liam Crowley, Douglas H. Boyes, Owen T. Lewis, Peter W.H. Holland

**Affiliations:** Department of Biology, University of Oxford, 11a Mansfield Road, Oxford OX1 3SZ, UK; UK Centre for Ecology & Hydrology, Wallingford, OX10 8BB, UK

## Abstract

Colour vision in insects is determined by signalling cascades, central to which are opsin proteins, resulting in sensitivity to light at different wavelengths. In certain insect groups, lineage specific evolution of opsin genes, in terms of copy number, shifts in expression patterns, and functional amino acid substitutions, has resulted in changes in colour vision with subsequent behavioural and niche adaptations. Lepidoptera are a fascinating model to address whether evolutionary change in opsin content and sequence evolution are associated with changes in vision phenotype. Until recently, the lack of high quality genome data representing broad sampling across the lepidopteran phylogeny has greatly limited our ability to accurately address this question. Here, we annotate opsin genes in 219 lepidopteran genomes representing 33 families, reconstruct their evolutionary history, and analyse shifts in selective pressures and expression between genes and species. We discover 44 duplication events in opsin genes across ∼300 million years of lepidopteran evolution. While many duplication events are species or family specific, we find retention of an ancient long-wavelength sensitive (LW) opsin duplication derived by retrotransposition within the speciose superfamily Noctuoidea (in the families Nolidae, Erebidae, and Noctuidae). This conserved LW retrogene shows life stage specific expression suggesting visual sensitivities or other sensory functions specific to the early larval stage. This study provides a comprehensive order-wide view of opsin evolution across Lepidoptera, showcasing high rates of opsin duplications and changes in expression patterns.

## Introduction

Opsin genes belong to the G protein-coupled receptor multigene family; the encoded opsin proteins form a photosensitive complex through covalent binding to a retinal-based chromophore (Palczewski et al. 2000; Briscoe and Chittka 2003; Nilsson 2009). The spectral sensitivities of an organism is directly linked to its opsin gene content and expression, thus this gene family provides a useful framework to relate changes in the genotype to the evolution of visual adaptations and behaviours (Arikawa 2017; Hauser and Chang 2017; van der Kooi et al. 2021; Van Nynatten et al. 2021). Opsin gene duplication and divergence, and gene loss, are two mechanisms by which insects have evolved altered visual sensitivities to adapt to niche-specific light environments and external cues (Spaethe and Briscoe 2004; Frentiu et al. 2007; Briscoe 2008; Thoen et al. 2014; Futahashi et al. 2015; Chen et al. 2016; Feuda et al. 2016; Sharkey et al. 2017; Armisén et al. 2018; Almudi et al. 2020; Feuda et al. 2021; Sondhi et al. 2021; Guignard et al. 2022; McCulloch, Macias-Muñoz, Mortazavi, et al. 2022). However, other mechanisms are known to increase the diversity of opsin specificities resulting in variation in light sensing abilities, such as amino acid substitutions without gene duplication causing shifts in absorption spectra (Shichida and Matsuyama 2009; Wakakuwa et al. 2010; Hauser et al. 2014; Liénard et al. 2021; Sharkey et al. 2023), co-expression of certain opsin proteins in a single photoreceptor cell (Wakakuwa et al. 2004; Perry et al. 2016; Satoh et al. 2017; Macias-Muñoz et al. 2019; Ilić et al. 2022), differences in ommatidia structure of the compound eye (Lau and Gross 2007; Lau and Meyer-Rochow 2007), and paralog specific opsin gene expression (Arikawa 2003; Perry et al. 2016; Finkbeiner and Briscoe 2021; Roberts et al. 2023).

In Lepidoptera (butterflies and moths), colour vision is determined by rhabdomeric type r-opsins encoded by three genes, with each opsin having a different peak wavelength sensitivity (λmax): long-wavelength-sensitive opsins (LW opsin; λmax 500–600 nm) which can respond to green light, short-wavelength-sensitive opsins (Blue or SWS opsin; 400–500 nm) sensitive to blue light, and ultraviolet opsins (UV opsin; 300-400 nm) which respond to ultraviolet light (Briscoe and Chittka 2003; Stavenga and Arikawa 2006; Henze and Oakley 2015; Feuda et al. 2016; van der Kooi et al. 2021). In addition to this core set of opsin genes, Lepidoptera also possess Rh7 and c-opsin genes, thought not to be linked to vision but whose full function and phylogenetic distribution is unknown (Feuda et al. 2016). Several duplications of opsin genes have been observed in lepidopteran species. For example, many butterflies in the genus *Heliconius* can see from the ultraviolet to the red extremes of the light spectrum, owing to duplication of the UV opsin gene followed by amino acid changes driven by positive selection in the ancestor of the clade (Briscoe et al. 2010). This also resulted in sexually dimorphic UV colour vision, with females of certain species able to distinguish between different UV wavelengths (Finkbeiner and Briscoe 2021; Chakraborty et al. 2023). This specialisation in vision is thought to have co-evolved with wing coloration (Finkbeiner et al. 2017). Similarly, a Blue opsin gene duplication in the family Lycaenidae allowed evolution of a green-shifted Blue opsin paralog which, in combination with a red-shifted LW opsin gene, may allow finer wavelength discrimination, again coinciding with wing coloration evolution (Bernard and Remington 1991; Sison-Mangus et al. 2006; Sison-Mangus et al. 2008; Liénard et al. 2021). Such diversity in the mechanisms of opsin evolution has resulted in massive variation in colour vision, habitat adaptation, and feeding habits in this diverse order of insects (van der Kooi et al. 2021). Additionally, the number of transitions between diurnal and nocturnal behaviour within this order (Kawahara et al. 2018) mean that this group is an ideal system to relate changes in opsin content and evolution with transitions in lifestyle and behaviour.

Despite these interesting case studies, much remains unknown concerning opsin diversity across Lepidoptera. The paucity of species sampling is also confounded by the fact that, until recently, high quality genome assemblies for this order have been limited in number, compromising efforts to annotate opsin genes, assign orthology, and confirm true gene losses (Feuda et al. 2016; Sondhi et al. 2021). Here we examine newly generated, chromosome level assemblies for 219 lepidopteran species, representing the largest, most phylogenetically representative dataset used to date. Chromosome-level genome assemblies publicly released by the Darwin Tree of Life project (The Darwin Tree of Life Project Consortium 2022) allow for accurate identification and analysis of opsin genes in all species. We assessed the rate of opsin gene duplication and loss across the tree, revealing dynamic changes in opsin copy number. Although there is conservation of five core lepidopteran opsin genes, we also identify 44 distinct duplication events, many of which occurred in the stem lineage of certain lepidopteran clades, including Tortricidae, Microterigidae, Lycaenidae, and Noctuoidea. Sequencing and analysis of transcriptomic data from members of the Noctuoidea superfamily revealed life stage-specific expression of LW paralogs, suggesting subfunctionalization of expression domains. Finally, we also tested whether transitions from nocturnality to diurnality are associated with shifts in evolutionary rate within the visual opsin genes.

## Results

From a dataset of 219 lepidopteran genome sequences representing 33 families we constructed a species tree and annotated opsin genes (Figure 1A, Supplementary figure S1). The dense sampling of species represents much of the lepidopteran diversity and includes species from the early diverging Microterigidae family. The dataset includes 152 nocturnal, 56 diurnal species and 11 species with both night and day-flying behaviours (Supplementary figure S1). These recently generated chromosome level genome assemblies, most generated by the Darwin Tree of Life Project (The Darwin Tree of Life Project Consortium 2022), allowed for more accurate and comprehensive identification of opsin gene open reading frames than in earlier studies. We used a combination of BLAST (Altschul et al. 1990) and Exonerate (Slater and Birney 2005) in an iterative approach to annotate opsin genes in all sampled genomes (see Material and methods). We identified 1,279 opsin genes across 219 lepidopteran species (Figure 1A). Phylogenetic reconstruction of opsin genes using a Maximum Likelihood approach recovers the monophyly of all main opsin groups: four r-opsins (UV, Blue, LW, and Rh7) and one c-opsin (Figure 1B, Supplementary figure S2). The branching patterns within the gene tree, in combination with gene structure and surrounding gene synteny where required, were used to infer the phylogenetic node of origin for each opsin gene duplication event (Figure 1C).

**Figure 1.**
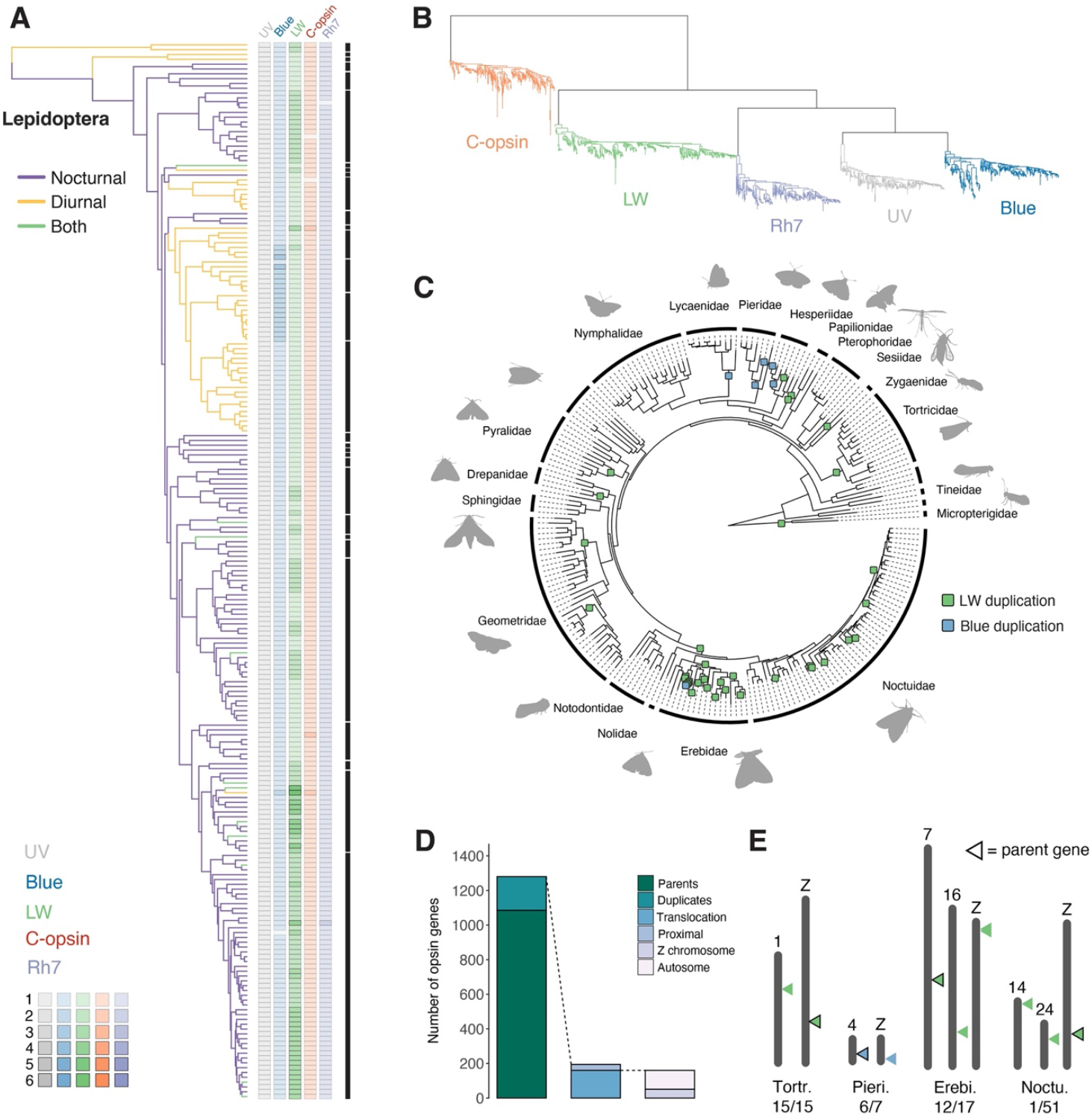
Opsin diversity and copy number across Lepidoptera. **(A)** Species phylogeny (left) with branch colours representing day/night flight activity of adults, along with copy number of each opsin gene type in each species (right). Each opsin gene has a distinct colour, with the strength of the colour representing copy number (ranges from 0-6 gene copies). Black bars represent lepidopteran families. Species names are given in Supplementary figure S1. **(B)** Gene tree of opsin genes present in Lepidoptera (amino acid sequences; maximum likelihood); each opsin gene groups into its own monophyletic clade. **(C)** Species tree with families labelled with black bars, and Blue and LW opsin gene duplication events mapped to their minimal branch of origin. **(D)** Bar chart representing the proportion of opsin genes which (i) underwent duplication in Lepidoptera, (ii) duplicated near the parent gene or were translocated to a new genomic location, and (iii) the number of translocation events resulting in duplicate copy placed on another autosomal chromosome versus sex chromosome (Z or W). **(E)** Opsin genes which translocated from an autosome to a sex chromosome mapped to chromosome graphs for each family where this occurred (Torticidae, Pieridae, Erebidae, and Noctuidae). Numbers show the number of species within the family which share this sex-linked opsin gene.

### Blue and LW opsin duplications were prevalent during lepidopteran evolution

We find variation in opsin gene copy number within different lineages, caused by 34 independent duplications of the LW opsin across 10 families, six duplication events of the Blue opsin in three different butterfly families and one species of moth, one Rh7 duplication, and three c-opsin duplications (Figure 1C). These include some previously described duplication events: the duplication of the Blue opsin in the butterfly family Lycaenidae (Bernard and Remington 1991; Sison-Mangus et al. 2006; Sison-Mangus et al. 2008; Liénard et al. 2021); a Blue opsin duplication within the Pieridae (Awata et al. 2009; Wakakuwa et al. 2010), which we find to be missing from the Wood white butterfly (*Leptidea sinapis*) and underwent an additional tandem duplication in the Clouded yellow butterfly (*Colias croceus*); and two LW duplications in the swallowtail butterfly *Papilio machaon* (Arikawa 2003; Saito et al. 2019). Increased taxon sampling in certain lineages refines the point of origin in lepidopteran evolution for certain duplication events. For example, the Blue duplication in Lycaenidae is present in all species in our dataset confirming that this event occurred in an ancestor to this family. Additionally, the Blue duplication in the Pieridae family is now placed on the branch after divergence from *Leptidea sinapis* (Figure 1C). We find another duplication of the Blue opsin within Hesperiidae, where a duplicate copy is shared between *Thymelicus sylvestris, Hesperia comma*, and *Ochlodes sylvanus*. We find it originated from a tandem duplication event in the ancestor of these species in Hesperiidae, and that there was also a further tandem duplication of the Blue opsin in *Ochlodes sylvanus* (Figure 1C). We also uncover a novel duplication of the Blue opsin in the day-flying Mother Shipton moth, *Euclidia mi* (Noctuoidea: Erebidae), which was the only example of duplication of this gene outside of the butterflies. This additional Blue opsin originated by tandem duplication from the parent opsin gene, and has transcriptional orientation in the opposite direction (Supplementary figure S3).

Duplication of the Blue opsin in butterflies has previously been shown to lead to shifts in wavelength sensitivities in the paralogous copies resulting in finer wavelength discrimination. In Lycaenidae, duplication of the Blue opsin resulted in a typical Blue opsin with λmax of 435 to 440 nm, and a green-shifted Blue opsin with λmax 495 to 500 nm (Liénard et al. 2021). In Pieridae, the duplication resulted in one normal Blue opsin and one violet-shifted Blue opsin with λmax of 420 nm (Wakakuwa et al. 2010). Spectral tuning sites responsible for these shifts in wavelength peak absorbance have been functionally characterised (Wakakuwa et al. 2010; Liénard et al. 2021), with a substitution from serine to alanine at position 116 responsible for a 5 to 13 nm shift in peak absorbance in both *Eumaeus atala* (Lycaenidae) and *Pieris rapae* (Pieridae). Two further substitutions (G175S and Y177F) combined to give a 73-nm bathochromic shift in *Eumaeus atala* (Liénard et al. 2021). We find the S116A substitution present in all paralogous Blue copies within Lycaenidae, implying this spectral shifting substitution occurred once following the duplication event (Supplementary figure S4). Within Pieridae, the S116A substitution is present in the closely related *Pieris rapae, Pieris napi*, and *Anthocharis cardamines* duplicated copies, however has reverted to alanine in *Pieris brassicae*. Within Hesperidae, all Blue opsin copies possess glycine at this site, while both Blue copies in *Euclidia mi* have the ancestral serine (Supplementary figure S4).

We discovered a large number of gene duplication events involving the LW opsin, amounting to 34 duplications across the species tree (Figure 1C). These include: a gene duplication shared between both Micropterigidae species; a duplication shared between all 15 species analysed in the family Tortricidae; two LW copies in the day-flying Six-spot burnet moth (*Zygaena filipendulae*) suggesting a recent duplication event; a gene duplication in the Small skipper butterfly *Thymelicus sylvestris*; an LW duplication on the branch leading to the Pyralidae, with a duplicate subsequently lost in two species in this clade; a shared duplicate in the two Drepanidae species; two independent duplications within the Geometridae; a shared duplicate within the Noctuoidea superfamily (Nolidae, Erebidae and Noctuidae families), as well as 23 subsequent duplication events of this paralog (see next Section) (Figure 1C). Duplications of the c-opsin gene were also found in the European swallowtail butterfly (*Papilio machaon*), the Mother Shipton moth (*Euclidia mi*), and the Buff-tip moth (*Phalera bucephala*) (Figure 1A).

Of the opsin genes present in greater than one copy (199 genes in 130 species), 163 of the duplicate genes (82%) are present on a different chromosome to their parent gene of origin (Figure 1D). This suggests a high rate of translocation following opsin duplication in Lepidoptera. There are 36 opsin genes which are in multi-copy and present on the same chromosome, either in tandem or proximal locations to the parent opsin gene. Many of these pertain to evolutionarily recent duplication events, such as the c-opsin duplicate in *Papilio machaon* and LW duplicates in *Spodoptera frugiperda, Laspeyria flexula*, and *Spilosoma lubricipeda*. Some opsin paralogs, present in all species in a family, such as the LW duplication in Micropterigidae and the Blue duplication in Lycaenidae (Figure 1C), are likely much older yet have been retained in close genomic association following radiation of the clade. In the case of the two Blue paralogs in all Lycaenidae species, these genes are tightly linked with an average of ∼6kb of intergenic sequence separating the two paralogs (10 species, range ∼3kb to ∼13kb). This suggests a selective pressure to retain close linkage.

Considering all opsin genes which underwent translocation following duplication, we find 51 opsin paralogs (from 36 species, representing 4 different taxonomic families) are located on the Z chromosome (Figure 1D & E); we do not find any opsin genes located on the female-specific W chromosome in this data set. This contrasts with a W-linked UV opsin gene in some *Heliconius* species giving female-specific expression in the heterogametic (ZW) sex (Chakraborty et al. 2023). We note, however, that only 59/219 of the genome assemblies in our dataset were constructed from female individuals, so there could be hidden diversity of opsin genes on the W chromosome in Lepidoptera. Nonetheless, the fact that many duplicated opsin genes are located on the homogametic sex chromosome (Z in Lepidoptera) gives potential for sex-specific regulation.

One example of translocation to the Z chromosome following gene duplication includes the LW opsins in the family Tortricidae (Figure 1C & E). All species within this family possess a duplicate LW copy on an autosome and the parent LW copy on the Z chromosome, which suggests relocation of the parent gene to a new genomic context following duplication on the ancestral Tortricidae branch. This gene translocation was the result of a fusion of the ancestral autosome to the Z chromosome in this family (Supplementary figure S5A) (Wright et al. 2023). It was possible to determine which LW gene was the ancestral, parent copy versus which was the recently duplicated copy due to the syntenic region (i.e. flanking genes) of the parent copy matching that of other species with a single LW opsin gene. This pattern is also seen in a Noctuid species (*Anorthoa munda*), where the parent LW copy is present on the Z chromosome, while the duplicate copies are on separate autosomes (Supplementary figure S5B). Conversely, the Blue paralog resulting from duplication within the family Pieridae, present in 6 out of 7 species sampled in our dataset (Figure 1C) is located on the Z chromosome in each of the species. This translocation is not a result of autosomal-chromosome fusion (Wright et al. 2023), but suggests that this gene was translocated to a new genomic location following duplication. Interestingly, this translocation to the Z chromosome resulted in the duplicate Blue opsin being located downstream and in the opposite orientations as a gene homologous to *paraplegin* (part of the AAA family proteins), and in the same orientation as a gene which contains a 3’5’-cyclic nucleotide phosphodiesterase, catalytic domain, in all species except for *Aporia crataegi*. Direct role of cyclic nucleotide phosphodiesterases in phototransduction remains uncertain, however, they have been shown to localise in photoreceptor cells in the fly *Calliphora erythrocephala* (Schraermeyer et al. 1993). In the family Erebidae, the LW opsin paralog located on the Z chromosome in 12/17 species was also likely due to an ancestral gene translocation event to the sex chromosome on the branch following the split from the subfamily Lymantriinae.

### An ancient, conserved LW retrocopy within the Noctuoidea superfamily shows lifestage specific expression

One of the most ancient cases of opsin duplication within the Lepidoptera within the Noctuoidea superfamily (at the base of the Nolidae, Erebidae, and Noctuidae families), where the LW opsin underwent duplication (Figure 1). Previously described in an erebid moth (Feuda et al. 2016) and a noctuid moth (Xu et al. 2016), we now suggest that this gene likely duplicated once, shared between three families within the superfamily Noctuoidea, by assessing the LW gene tree (Figure 1B, Figure 2A, Supplementary figure S6), which shows two distinct monophyletic LW opsin groups, here named LWS1 and LWS2 (Figure 2A). Given that this superfamily diverged ∼80 million years ago (Kawahara et al. 2019), it is particularly striking that the two copies are retained in every species in our dataset, consistent with functional divergence and selective retention. Further evidence for the potential adaptive benefit of this ancient LW duplication is the finding that further duplication events of this LWS2 gene occurred on 19 separate branches within Erebidae and Noctuidae, 13 of which are recent events (species specific in our dataset; Figure 1C). The LWS2 is intronless in all species (Feuda et al. 2016; Xu et al. 2016), compared to LWS1 which contains 7 introns, indicating that LWS2 is derived from LWS1 by retrotransposition (Betrán et al. 2002; Booth and Holland 2004; Kaessmann 2010). Further support for retrotransposition as the mechanism of origin is the fact that LWS2 is always located on a separate chromosome from LWS1.

**Figure 2.**
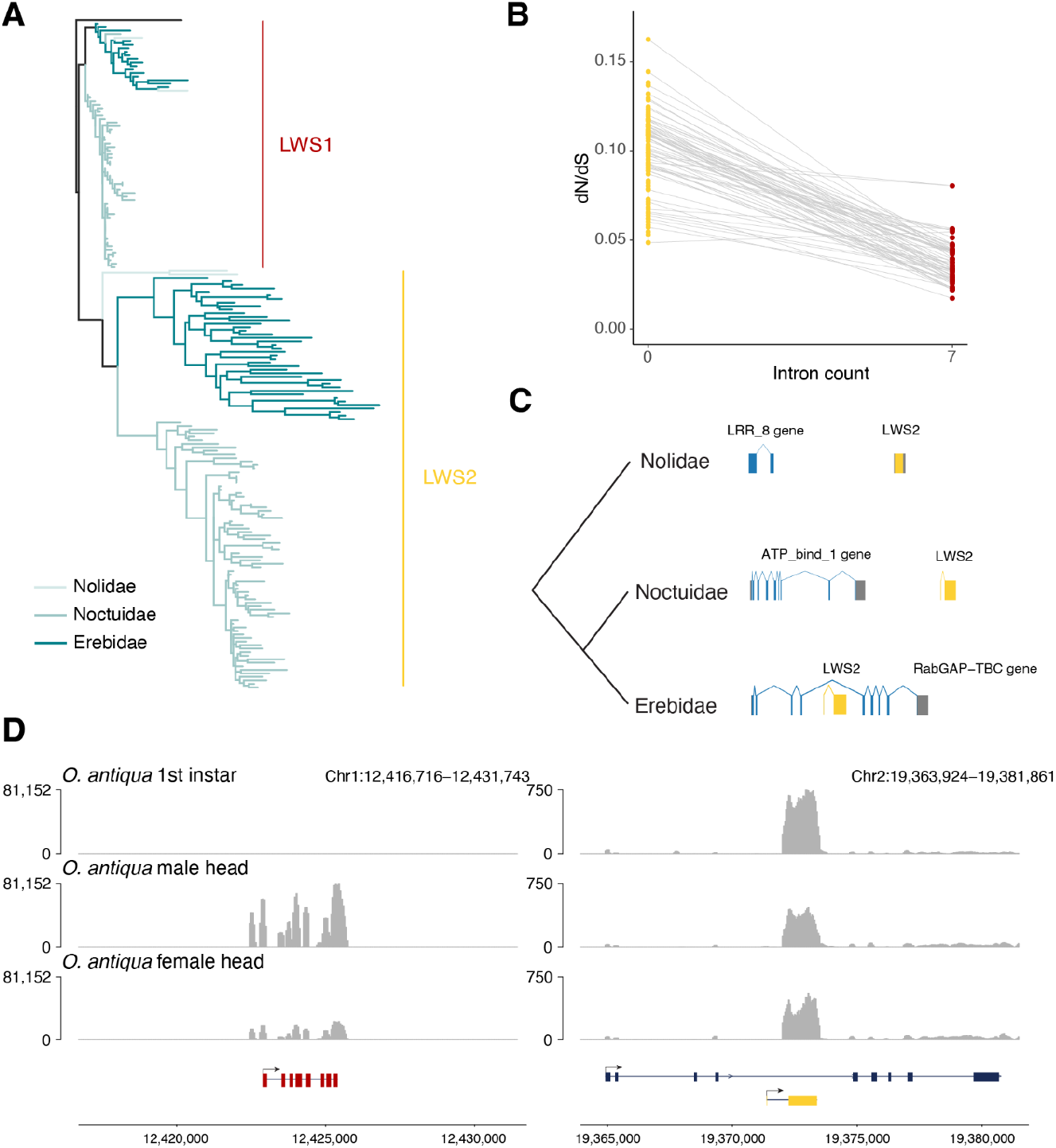
Evolution and expression of LW duplicate within Noctuoidea. **(A)** Gene tree of LW opsin genes in Noctuoidea families Nolidae, Erebidae, and Noctuidae, with branches coloured according to family. LWS1 (red) represents the ancestral parent LW opsin gene, LWS2 (yellow) represents LW paralog duplicated at the base of the Noctuoidea superfamily. **(B)** dN/dS values of both parent and duplicate LW copies in Noctuoidea, intronless copy represents the LW paralog while the parent LW opsin contains the normal 7 introns. Grey lines linking points (LW genes) represent parent-duplicate copies in a single species. **(C)** Syntenic locations of the LWS2 genes in Nolidae, Noctuidae, and Erebidae. LWS2 gene is shown in yellow, and the closest upstream gene in the same orientation is shown in blue. Blocks in genes represent exons, while lines joining them represent introns. **(D)** Expression levels of parent (left) and duplicated (right) LW copy in *Orgyia antiqua* (Erebidae). Tracks represent depth of mapped RNA reads on the genome. Top track shows expression levels for 1st instar larva whole body, second track shows male head, third track shows female head. Chromosome locations are given for both copies, and the gene structure (including orientation and exons/intron) are shown below.

Examining branch lengths in the gene tree of Noctuoidea LW opsins, it is clear that the protein encoded by intronless LWS2 genes is evolving at a faster rate than the protein encoded by the parent LWS1 gene (Figure 2A). Assessing dN/dS values, we find increased global dN/dS ratio in the LWS2 gene relative to its parent gene LWS1, suggesting relaxation of purifying selection following the duplication event (Figure 2B). This relaxation in selection was confirmed using a robust model-based approach, applying RELAX implemented in Hyphy (Wertheim et al. 2015; Kosakovsky Pond et al. 2020), which measures shifts in the stringency of selection acting on a gene; where a value of k>1 indicates intensified strength of selection, and k<1 indicates relaxation of selection strength. Measuring shifts (i) on the branch leading to the LWS2 clade (k = 0.63, *p* = 0.02) and (ii) on all branches within the LWS2 clade of the gene tree (k = 0.334, *p* = 0), we find significant relaxation in the intensity of selection acting on the duplicate LWS2 gene relative to the background rate of the LWS1 clade (Figure 2A). We next tested for positive selection in both LW copies in Noctuoidea using the branch-site model aBSREL implemented in HyPhy (Smith et al. 2015). Testing the branch leading to the LWS2 clade, and separately all branches within the LWS2 clade, we cannot identify amino acid sites with significant evidence for positive selection, relative to the background of LWS1. This suggests that, while there is relaxation of selection on the LWS2 copy, there is still overall selective pressure maintaining the function of this gene in all species.

The genomic location of the intronless LWS2 duplicate copy differs between taxonomic families within Noctuoidea (Nolidae, Erebidae, and Noctuidae; Figure 2C, Supplementary figure S7). In the two Nolidae species (*Meganola albula* and *Nycteola revayana*) the LWS2 retrogene is present in conserved syntenic regions, located downstream and in the same orientation as a Leucine rich repeat domain-containing gene (LRR-8), homologous to *Connectin* in *Drosophila melanogaster* (Figure 2C, Supplementary figure S7). In the 17 Erebidae species, the LWS2 gene is present in a different genomic location, within the intron of another gene (a Rab GTPase gene) in all species (Figure 2C, 2D right, Supplementary figure S7 & S8). As noted above, this LWS2 locus underwent an additional duplication and subsequent translocation in some Erebidae species (Figure 1C), with translocation resulting in LWS2 copies on the Z chromosome of 12 species in our dataset. Of these, 8 have multiple copies of LWS2 on the Z chromosome suggesting further tandem duplication following the translocation event. In the 51 Noctuidae species, the LWS2 copy is always present in the same syntenic region in all species in the family, downstream and in the same orientation of an ATP bind1 domain containing gene (Figure 2C, Supplementary figure S7).

It was previously observed that the intronless LWS2 duplicate gene was more highly expressed in the first instar larval stage of the noctuid moth, *Helicoverpa armigera*, while the parental LWS1 gene has higher expression in the adult (Xu et al. 2016). To assess the consistency of this pattern, we measured the expression level of all opsin genes in larval and adult stages in an Erebidae species, *Orgyia antiqua* (Vapourer moth). RNA extraction was carried out on 1st instar larvae of *Orgyia antiqua*, as well as heads of adult male and adult female *Orgyia anitqua* (see Materials and methods). We found that the LWS1 gene is highly expressed in the adult stage while the duplicate LWS2 gene is most highly expressed in the 1st instar larval stage (Figure 2D, Supplementary figure S9). This conservation of expression domains, combined with the selective pressure analyses, suggests functional importance of the LWS2 gene in early larval stages of erebid and noctuid moths.

### Evidence of divergent molecular evolution in opsins of day-flying species

To investigate the patterns of molecular evolution of opsin genes relative to behaviour and lifestyle of the lepidopteran species we carried out tests to measure the strength and form of selection acting on each gene. First, we classified each species in our dataset based on whether the adults were nocturnal (152 species), diurnal (56), or have evidence for being both day and night-flying (11) (Figure 1A, Supplementary figure S1). For each of the opsin genes with functions directly related to vision (UV, Blue, and LW) we measured rates of synonymous (dS) and non-synonymous substitutions (dN), as well as their ratio (omega = dN/dS), first using a codon model where omega does not vary across site or branches in the tree (Muse and Gaut 1994). Comparing these values between the three lifestyle classes, we find that diurnal species have significantly higher dN/dS values in each visual opsin gene compared to the other two lifestyle categories (UV; *p* = 0.04146, Blue; *p* = 3.591e-06, LW; *p* = 0.0001309) (Figure 3A), suggesting different selective pressures acting on the opsin genes in diurnal lepidopteran species.

**Figure 3.**
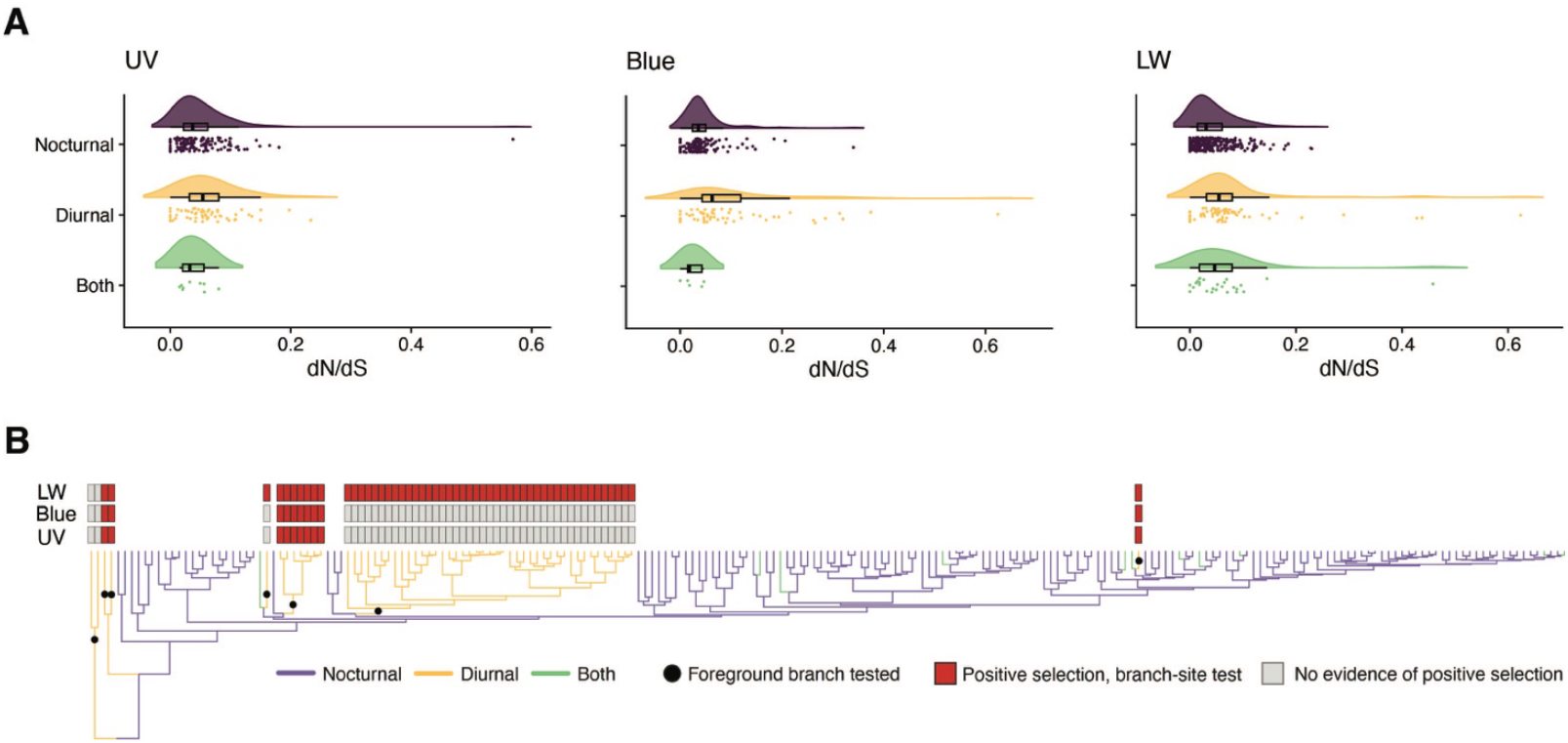
Levels of selection acting on opsin genes in species with different photic niches. **(A)** Level of selective pressure (dN/dS) acting on opsin genes in species with nocturnal activity (purple), diurnal activity (yellow), or evidence for both (green). Raincloud plots show dN/dS values for all opsin genes in each species in the dataset. (B) Results from branch-site test on the three opsins. Species tree below has branches colour by photic niche of each species, and black circles represent foreground branches tested for evidence of positive selection. Grey bars indicate no positive selection found in tested foreground branches, while red bars show opsin genes with some evidence of positive selection in the foreground branches.

To investigate further, we then tested for positive selection acting on all branches within the clades containing day-flying species in the orthologous opsin genes using the BUSTED-PH model in HyPhy (Kosakovsky Pond et al. 2020). This tests for episodic diversifying selection on foreground branches with the same trait where there is no evidence of positive selection on the background branches. For the Blue and LW opsins, we find evidence that selective pressures are significantly different between the background (nocturnal) and foreground (diurnal) branches; however, there is evidence for selection on both sets of branches. The UV opsin showed no evidence of episodic diversifying selection in the tested branches. Finally, to test for positive selection acting on specific sites we applied both the random sites model and the branch-site model implemented in PAML (Yang 2007) to the UV, Blue, and LW genes, with transitional branches to diurnal species labelled as foreground. We tested seven independent transitions from nocturnal to diurnal species (Figure 3B, Supplementary figure S1). The branch-site model revealed positive selection in all three opsin genes, with the LW opsin in particular having the highest number of sites with evidence for change due to positive selection, with 31 unique sites in total under positive selection in all branches tested (Figure 3B, Supplementary figure S10). In the case of the LW and Blue opsins, only one of the positively selected sites overlaps with known retinal binding sites and may directly affect spectral tuning (Supplementary figure S10). There are positively selected sites in all three of the tested opsins that map to the transmembrane regions near the retinal binding sites (Supplementary figure S10), which may have indirect impact on opsin spectral sensitivity. Further experimental work will be required to test whether these impact opsin spectral tuning.

While we find evidence that rates of sequence evolution are different in day-flying species, a phylogenetic correlation test did not find any evidence that gene duplication of the visual opsin genes was associated with diel-niche (Pagel’s λ; p = 0.66). This suggests that opsin duplication alone does not predict photic niche in a species, but may have a wider range of implications, such as sub-functionalization of expression or changes in sensory roles outside of vision.

## Discussion

The Lepidoptera provide a particularly interesting group to study the evolution of vision-related genes given the changes in behaviour through the life-cycle and the number of independent transitions from nocturnal to diurnal activity of adults. Colour discrimination is also known to differ between families and, in some cases, even between closely related species or sexes of the same species. Shifts in sensitivities to different wavelengths can occur through a range of different mechanisms, including structural changes to ommatidia, changes to opsin expression in specific photoreceptor cells, and processes of molecular evolution such as spectral-shifting amino acid substitutions in opsin genes. The role of opsin gene duplication and loss is also relevant and has been studied across arthropods, revealing some dramatic cases of gene turnover (Futahashi et al. 2015; Sharkey et al. 2017; Almudi et al. 2020; McCulloch, Macias-Muñoz, and Briscoe 2022). Recent advances in characterising the spectral sensitivity of opsin proteins in vitro provides a promising platform to pinpoint the functional role of opsin duplication and loss in driving variation in visual sensitivities (Liénard et al. 2021; Liénard et al. 2022).

A major question in the field is whether there is a relation between diel activity and opsin gene evolution. Here, we exploited newly generated high quality genome assemblies to compare the opsin genes between 219 species of Lepidoptera, representing a broad phylogenetic span. Overall we find a conserved opsin gene complement across Lepidoptera, with only one putative loss in the visual opsin genes: loss of the Blue opsin in the Twin-spotted Quaker moth (*Anorthoa munda*). We uncover 44 cases of duplication in the opsin genes, 40 of which affect the Blue (short wave sensitive) and LW (long wave sensitive) visual opsin genes, but no cases of duplication of the UV opsin. Amongst the duplicated genes, we find four cases of translocation of an opsin paralog to the Z chromosome following duplication. These events have the potential to lead to sex-linked expression differences related to dosage effects between homogametic males (ZZ) and heterogametic females (ZW). Whether such potential is realised may be dependent on the mechanism of transposition: either by chromosome fusion or movement of a small genomic region. Following autosome fusion to an ancestral Z chromosome, the distinct regions of the composite lepidopteran sex chromosome retain their respective patterns of dosage compensation, with the (now transferred) neo-Z region having a two-fold increase in transcription in ZZ females (Gu et al. 2019). We suggest this may be the case for the Tortricidae LW gene duplicate, which we find was translocated to the Z chromosome through autosomal fusion in the ancestor of this family. In contrast, the translocation of the Blue opsin duplicate to the Z chromosome in Pieridae was not a result of chromosomal fusion but rather a translocation of a DNA locus; we suggest this newly Z-linked opsin is likely to have dosage balance between sexes. It is also notable that this single gene translocation resulted in the Blue paralog located downstream of a 3’5’-cyclic nucleotide phosphodiesterase gene. The orthologous gene in *Drosophila melanogaster* is expressed in photoreceptor cells, and thus this locus may share regulatory sequences with the translocated opsin gene.

Of the 44 opsin duplication events identified, 16 occurred within lineages containing day-flying species. The majority of Lepidoptera are nocturnal with day-flying being a derived trait in most, if not all cases. Opsin duplication in day-flying species shows some interesting lineage specific patterns, such as an LW duplication shared between the Micropterigidae species, LW duplication in the Six-spot burnet moth (*Zygaena filipendulae*), Blue opsin duplication in the Mother Shipton moth (*Euclidia mi*), and five independent duplications of the Blue opsin within the butterflies. These duplications could provide a novel source of variation to generate shifts in visual spectra for these species (Wakakuwa et al. 2010; Liénard et al. 2021). However, while these are interesting individual candidates for opsin duplication related to diurnality, we did not find a significant correlation between opsin duplication and diel activity across our species phylogeny (Pagel’s λ; *p*=0.66), suggesting that opsin duplicates may have been co-opted for a range of possible roles relating to vision and other sensory functions (Feuda et al. 2022). One such case of opsin duplication not directly related to diurnal behaviour is an ancient LW duplication which likely occurred at the base of three families (Nolidae, Erebidae, and Noctuidae) within the superfamily Noctuoidea. This event, which occurred via retrotransposition (all copies possess no introns), generated a new LW opsin paralog, LWS2, which was then retained in every species in our dataset. Differential expression of the two LW copies between life stages has now been shown in two species from different families (Noctuidae and Erebidae) and likely represents the ancestral state, although more data for species in the Nolidae family is required to confirm this. This route to subfunctionalization of opsin paralogs through changes in temporal or spatial expression has been found in other insects, for example in dragonflies, mayflies, mosquitos, and other lepidopterans (Futahashi et al. 2015; Giraldo-Calderón et al. 2017; Almudi et al. 2020; Kuwalekar et al. 2022). In many of these other cases the larval and adult stages occupy very different niches and light environments (larvae are aquatic while adults are terrestrial). This is not the case for moth species within the Noctuoidea superfamily (most larvae and adults are nocturnal). Interestingly, we note that early instar larvae of Noctuidae and Erebidae are highly active and have rapid dispersal while some other larvae show more sedentary behaviour in early instars. Whether the function of the LWS2 retrocopy is linked to this behaviour is unknown.

While the exact function of LWS2 requires further analyses, it highlights the role of retrotransposition in generating new gene copies, with the potential for different functions (Kaessmann 2010; Carelli et al. 2016). Retrotransposition of opsin genes has also been noted in Diptera, where the *Rh3* gene (UV sensitive opsin) and *Rh6* gene (LW sensitive) originated via this mechanism in the *Drosophila* genus and different mosquito lineages, respectively (Giraldo-Calderón et al. 2017; Feuda et al. 2021). Retrogenes are easy to distinguish from their parent copy due to the missing introns, but importantly from an evolutionary perspective will not inherit promoters or most regulatory elements. This suggests that shifts in expression patterns are likely and could be driven by genomic features around the new insertion site in the genome (Carelli et al. 2016). In the case of the LWS2 retrogene, it is unclear where the original insertion site was because the retrogene has a different genomic location in the three extant families examined. In Erebidae, LWS2 is located within the intron of another gene (Figure 2C, Supplementary figure S8), a RabGAP-TBC domain-containing gene that we propose is homologous to human TBC1 domain family member 20 (TBC1D20). Interestingly, TBC1D20, along with five other TBC domain proteins, are crucial for trafficking G-protein-coupled receptors (GPCRs), to which opsins genes belong, from the endoplasmic reticulum, through the Golgi apparatus, to their final location in the plasma membrane (Wei et al. 2019). Integration into the intron of this host gene may have provided crucial transcriptional as well as functional integrity to the duplicate LW copy within Erebidae. We note, however, that while LWS2 is expressed strongly in early instar larvae of the Vapourer moth *Orgyia antiqua*, we find relatively low expression of the TBC1D20 gene (Figure 2D).

## Conclusion

We uncover extensive and previously underappreciated opsin gene duplication and evolutionary change in the Lepidoptera. Due to broad sampling of high quality genomes, we could confirm gene losses and the timing of duplication events using gene structure and genome synteny. We find different modes of gene duplication, including retrotransposition and tandem duplication, both sometimes followed by translocation or chromosomal rearrangement. These events have provided opportunity for substitution accumulation and sequence divergence which has likely increased the transcriptional and functional diversity of opsin genes in this group (Lynch and Conery 2000; Kaessmann 2010). For example, we find evidence for transcriptional divergence between LW opsin genes in different lifecycle stages of some moths, and evidence for differential selective pressures acting on the opsin genes of day-flying moths and butterflies.

## Materials and Methods

### Data acquisition

The majority of the genomes used in this analysis were produced by the Darwin Tree of Life project (The Darwin Tree of Life Project Consortium 2022) which can be found on the Darwin Tree of Life (DToL) portal page (https://portal.darwintreeoflife.org) or under accession number PRJEB40665 in the European Nucleotide Archive (ENA; https://www.ebi.ac.uk/ena/browser/home). The genomes from the remaining additional lepidopteran species were obtained from NCBI. A list of all species and their associated genomes and sources can be found in Supplementary table S1.

### Species tree reconstruction

In order to map the presence, absence and copy number of the opsin genes to the species we required a species tree. Species tree reconstruction was carried out using a dataset of 1,465 genes annotated with BUSCO v5.1.2 (Manni et al. 2021) using the Lepidoptera gene set. Each gene annotated this way contained all species sampled and were aligned with MAFFT v7.4 (Katoh et al. 2005) and trimmed with trimAl (Capella-Gutiérrez et al. 2009) before concatenation into a supermatrix using the create_concatenation_matrix option in PhyKIT (Steenwyk et al. 2021). The species tree was inferred from this supermatrix using IQ-TREE v2.0 (Minh et al. 2020) and the LG model with a gamma distribution with four categories. Tree visualisation was carried out using Toytree (Eaton 2020) and ggtree (Yu et al. 2017).

### Opsin gene annotation

A total of 615 protein sequences obtained from two studies on opsin evolution in Diptera (Feuda et al. 2021) and Lepidoptera (Sondhi et al. 2021) were used as seeds in an initial tBLASTn search of the lepidopteran genomes used in this analysis. Open reading frames (ORF) of opsins discovered by the BLAST search were constructed using Exonerate v2.4 (Slater and Birney 2005). Next, to ensure accurate annotation of the full ORF for each opsin gene, we reran the tBLASTn and exonerate pipeline this time using the newly found opsin genes which had a start codon and appropriate protein length. Opsin genes were assigned to a specific type (UV, Blue, LWS, c-opsin, RH7) based on our initial BLAST and exonerate annotation, as well as a subsequent BLASTp search against the nr BLAST database. Finally, accurate classification of the opsin genes was confirmed by building a gene tree of all opsins (see next section) and ensuring each annotated opsin grouped in the correct clade in the tree.

### Opsin gene tree inference

Inference of the opsin gene tree was important for a number of parts of this study. All inferred opsin genes were aligned using MAFFT v7.4 (Katoh et al. 2005). To identify the relationships between the opsin genes we performed tree inference using Maximum Likelihood with IQ-Tree (Minh et al. 2020) applying ModelFinder to find the model of best fit (Kalyaanamoorthy et al. 2017). The gene trees were used to confirm correct annotation of the opsin genes by the sequence homology pipeline described above. Additionally, the patterns in the complete opsin gene tree were used as one line of evidence to infer when duplication events occurred along the species tree, by carrying out a manual inspection of the gene tree to reconcile the gene tree with the species tree. Finally, individual gene trees were also constructed for each of the opsin types using the same approach as above, and these were used when carrying out the tests for selective pressure (see next section).

### Selective pressure analyses

Tests for selection were carried out on the opsin genes in a number of different analyses. First, regarding the LW duplication event in Noctuoidea, we calculated rates of synonymous substitutions (dS) and nonsynonymous substitution (dN) per site, as well as their ratio (dN/dS), for both copies of this gene (i.e. LWS1 and LWS2 as described in the Results section). First, protein sequences for all copies of LWS1 and LWS2 in Noctuoidea species were aligned using MAFFT v7.4 (Katoh et al. 2005). Next, codon alignments were created using the protein alignments and corresponding nucleotide sequences as input to PAL2NAL (Suyama et al. 2006). dN/dS was calculated for all copies of the Noctuoidea LW genes using the Muse-Gaut (MG94) model (Muse and Gaut 1994) (--type local) implemented in HyPhy (Kosakovsky Pond et al. 2020) and the Noctuoidea LWS gene tree (Figure 2A). We also carried out a test for signatures of relaxation or intensification of selection, with the branches leading to and within the LWS2 clade set as the foreground and the LWS1 clade set as the background. To test this we employed the RELAX model (Wertheim et al. 2015) in HyPhy (Kosakovsky Pond et al. 2020). Finally, the aBSREL model in HyPhy (Kosakovsky Pond et al. 2020) was employed to test for evidence of positive selection on any branches in the Noctuoidea LW gene tree. This model, which does not require a priori partitioning or selection of branches on the phylogeny, estimates dN/dS on all branches of the tree (Kosakovsky Pond et al. 2011).

We used several tests to measure selection within the day-flying lineages relative to nocturnal species in our dataset. First, we calculated the dN/dS for each opsin gene in each species using the Muse-Gaut (MG94) model in HyPhy (Kosakovsky Pond et al. 2020) (--type local). Each dN/dS value for every species was used to summarise the general patterns of selection between day-flying species, night-flying species, and species with evidence of both. These summarised group values were visualised using Raincloud plots in R (Allen et al. 2019). Next, we used the BUSTED-PH test within HyPhy (Kosakovsky Pond et al. 2020) with all day-flying branches in the tree labelled as foreground test branches. Finally, we used codon based models employed in codeml (Yang and Nielsen 2002) using the pipeline Vespasian (zenodo.org/record/5779869; github.com/bede/vespasian), to test for evidence of episodic events of positive and divergent selection on selected branches leading to day-flying lineages in the lepidopteran phylogeny. We employed both site and branch-site models, comparing standard nested models using likelihood ratio tests, as implemented in Vespasian. Sites found to be under positive selection were mapped to protein models by predicting transmembrane helices for all three opsins using Phobius (Käll et al. 2007) through the webserver Protter (Omasits et al. 2014). Retinal binding sites were inferred by including data from the jumping spider rhodopsin-1 into the model (Varma et al. 2019).

### RNA sequencing and opsin expression quantification

A captive-reared female *Orgyia antiqua* was mated with a wild caught male; resultant fertilised eggs were maintained over winter at ambient temperature (UK). On the day of larval hatching, 30 first instar larvae were homogenized using a sterile needle in RNAProtect; total RNA was purified several days later using a QIAGEN RNAeasy Plus Micro kit. Eggs from the same mating were reared until adult emergence the same year; one male and one female head were processed for RNA extraction directly using an QIAGEN RNAeasy Plus Micro kit, after removal of antennae. Paired-end 150 bp Illumina RNA-seq was performed commercially by Novogene (www.novogene.com) using poly-A selection, random hexamer priming and amplification giving 5.5 to 7.9 Gb of sequence per sample (NCBI SRA accessions: SRX20677428, SRX20677423, SRX20677422).

RNA-seq data from the three samples (first instar larvae, male head, female head) were trimmed for quality using Trimmomatic v0.39 (Bolger et al. 2014). Next, transcriptome assembly was performed for each sample using Trinity v2.8.5 (Grabherr et al. 2011), and transcript abundance was calculated using kallisto v0.44 (Bray et al. 2016). Each opsin gene was identified by performing a reciprocal BLAST search, and gene expression abundance was measured using transcripts per million (TPM). Additionally, processed RNA reads were mapped to the *Orgyia antiqua* genome using bowtie2 (Langmead and Salzberg 2012) in order to visualise RNA read depth and expression of the LW opsin genes. Read coverage was quantified using bedtools v2.25.0 (Quinlan and Hall 2010), and gene tracks and RNA coverage were visualised using trackplot; github.com/PoisonAlien/trackplot (Pohl and Beato 2014).

## Acknowledgements

We thank Thomas Lewin for constructive discussions and advice. We also acknowledge the incredible effort of all members associated with the Darwin Tree of Life project, without whom this research would not be possible. This research was funded by the Wellcome Trust Darwin Tree of Life Discretionary Award (218328) and the John Fell OUP Research Fund. Finally, this research began as a direct influence of the multidisciplinary nature of Douglas Boyes, to whom we will always be grateful.

## Data availability

All data and code required to reproduce analyses and figures can be found in the Supplemental Materials and at GitHub (github.com/PeterMulhair/Lepidoptera_opsins).

## Notes

https://github.com/PeterMulhair/Lepidoptera_opsins

### Competing Interest Statement

The authors have declared no competing interest.

