## Supplementary file for "Opsin gene duplication in Lepidoptera: retrotransposition, sex linkage, and gene expression"

#### Supplementary Figures

Supplementary Figure S1: Lepidoptera species tree.
Supplementary Figure S2: Opsin gene tree.
Supplementary Figure S3: *Euclidia mi* Blue opsin tandem duplication.
Supplementary Figure S4: Gene tree and alignment of blue opsin duplicate genes.
Supplementary Figure S5: Autosome-Z chromosome fusion events in Tortricidae and *Anorthoa munda*.
Supplementary Figure S6: Noctuoidea LW opsin gene tree.
Supplementary Figure S7: Summary of LWS2 paralog genomic locations and duplication history.
Supplementary Figure S8: Genomic locations of LWS2 paralog in Erebidae species.
Supplementary Figure S9: TPM values for opsin genes in different lifestages of Vapourer moth (*Orgyia antiqua*).
Supplementary Figure S10: Opsin structure with retinal binding sites and overlap with posi- tively selected sites.

#### Supplementary Tables

Supplementary Table S1: List of species and their accession numbers used in this study.

#### Supplementary Figures

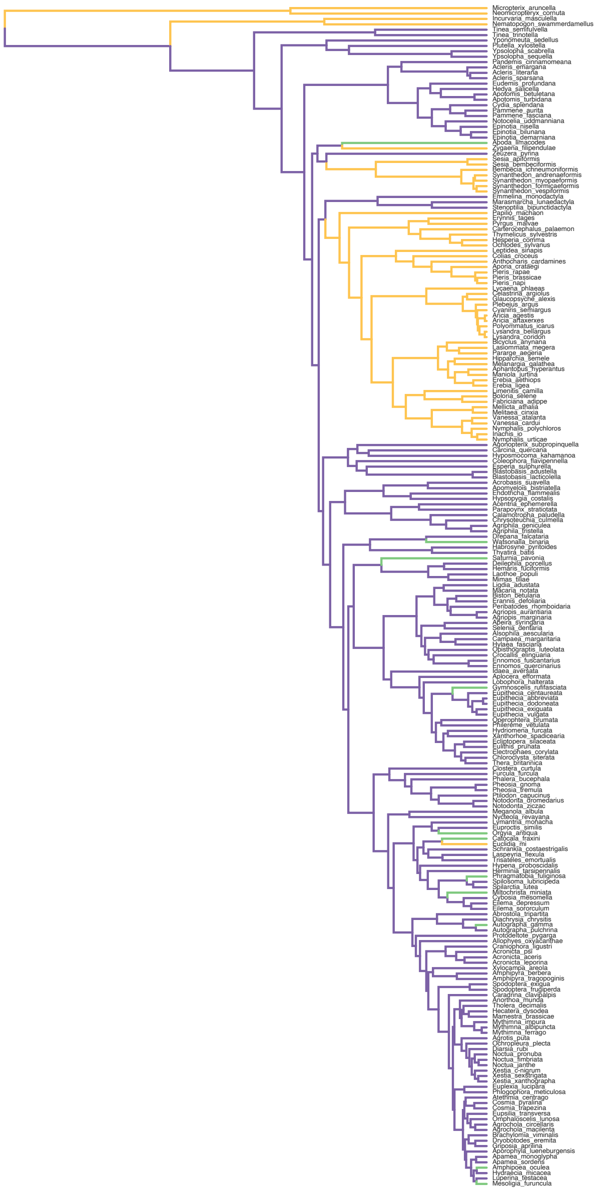

**Supplemental Figure S1: Lepidoptera species tree.** Species tree for representative Lepidoptera species inferred from BUSCO gene set of 1,281 genes, using a maximum likelihood supermatrix approach. Species names are given, and their corresponding branch colour denotes whether they are nocturnal (purple), diurnal (yellow), or have evidence for activity in both (green). The tree and branch colours correspond to those in the species tree in Figure 1A in the main text.

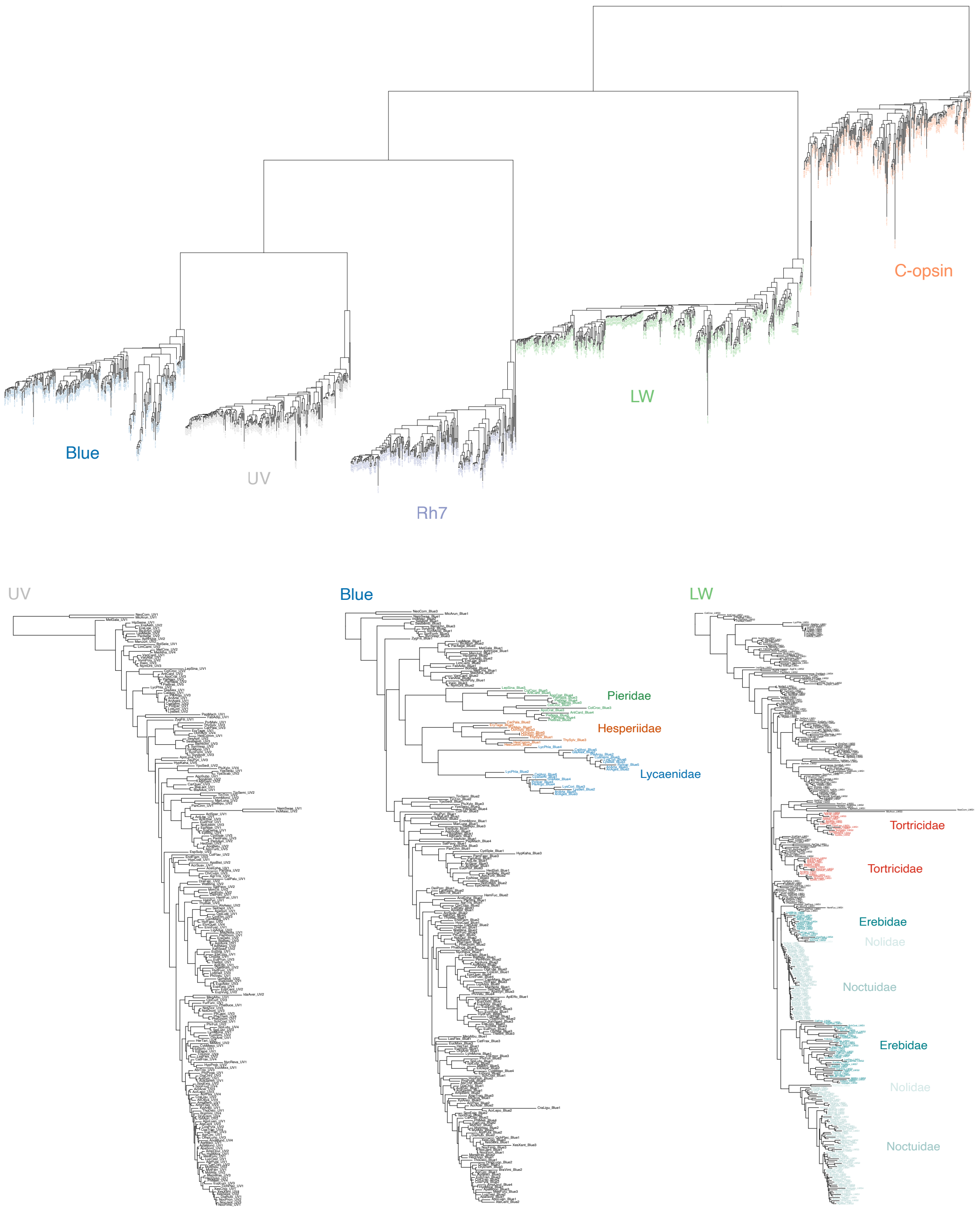


**Supplemental Figure S2: Opsin gene trees.** (Upper) Gene tree of all opsin sequences inferred using a maximum likelihood approach using IQtree. The tree and gene clade colours correspond to those in Figure 1B in the main text. (Lower) Gene trees for each of the visual opsins (UV, Blue, LW), with major duplications shared between a large number of species coloured and labelled with the lepidopteran family within which it occurred.

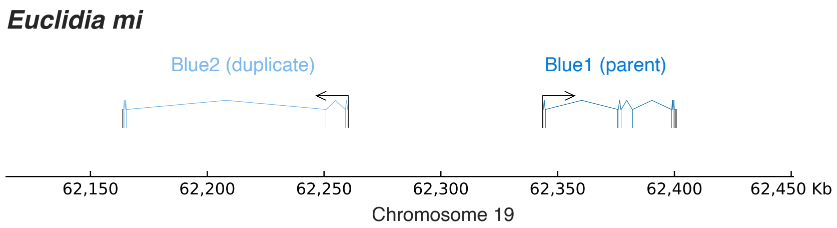


**Supplemental Figure S3: *Euclidia mi* Blue opsin tandem duplication.** Gene tracks showing tandem duplication of the Blue opsin gene in *Euclidia mi* (Mother Shipton moth). Both blue copies are located on chromosome 19, with the duplicated Blue opsin located 83kb upstream of the parent copy in the opposite orientation

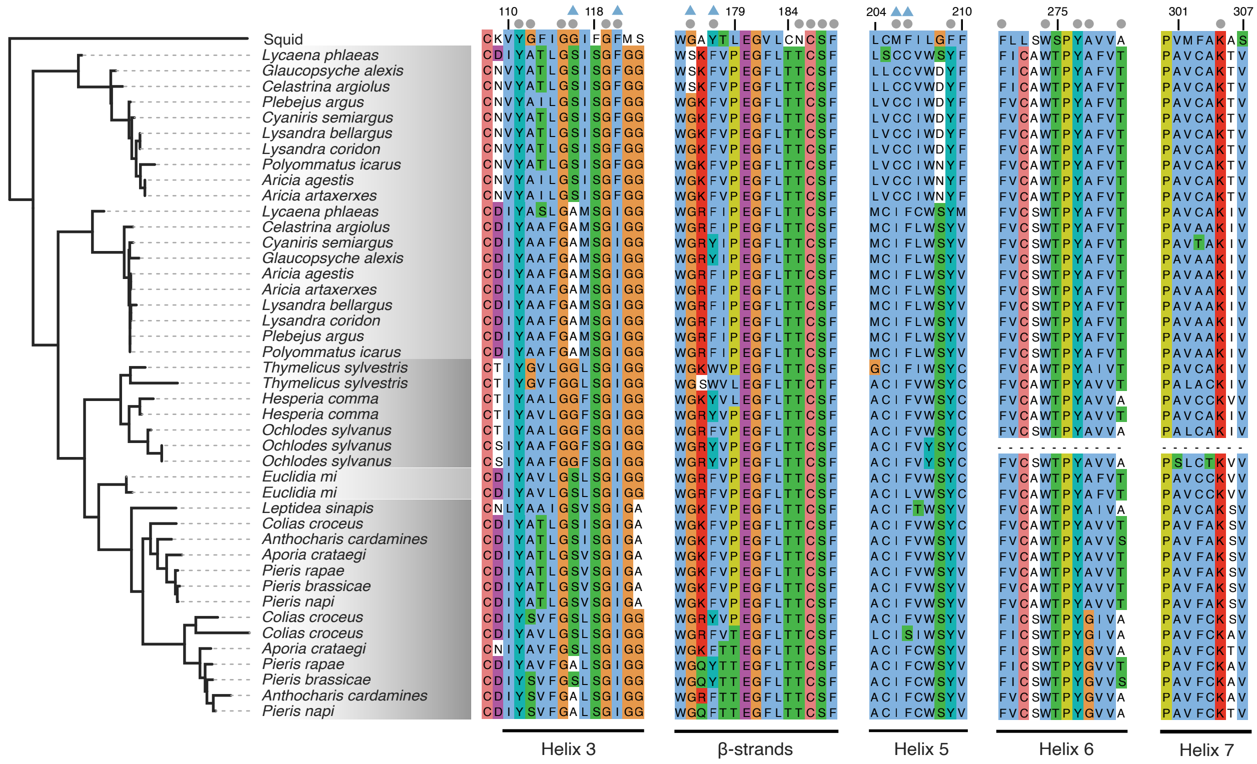


**Supplemental Figure S4: Gene tree and alignment of blue opsin duplicate genes. Left;** Maximum likelihood gene tree of blue opsin genes which underwent duplication in certain lepidopteran species. Alternating grey colours correspond to specific families within which the blue opsin gene underwent a duplication event (Lycaenidae, Hesperidae, Erebidae, and Pieridae). The Squid opsin (taken from *Todarodes pacificus*, accession no. CAA4990) is used as an outgroup. **Right;** Partial alignment of the corresponding blue opsin paralogs, showing regions of the opsin protein corresponding to helix loops and beta-strands. Dots and triangles above the alignment are obtained from (Liénard et al. 2021) and correspond to sites within 5 Å of any carbon atom in the retinal polyenechain, and sites shown to result in spectral shift of the blue opsin gene, respectively. The numbered sites above the alignment correspond to amino acid sites within the squid opsin protein.

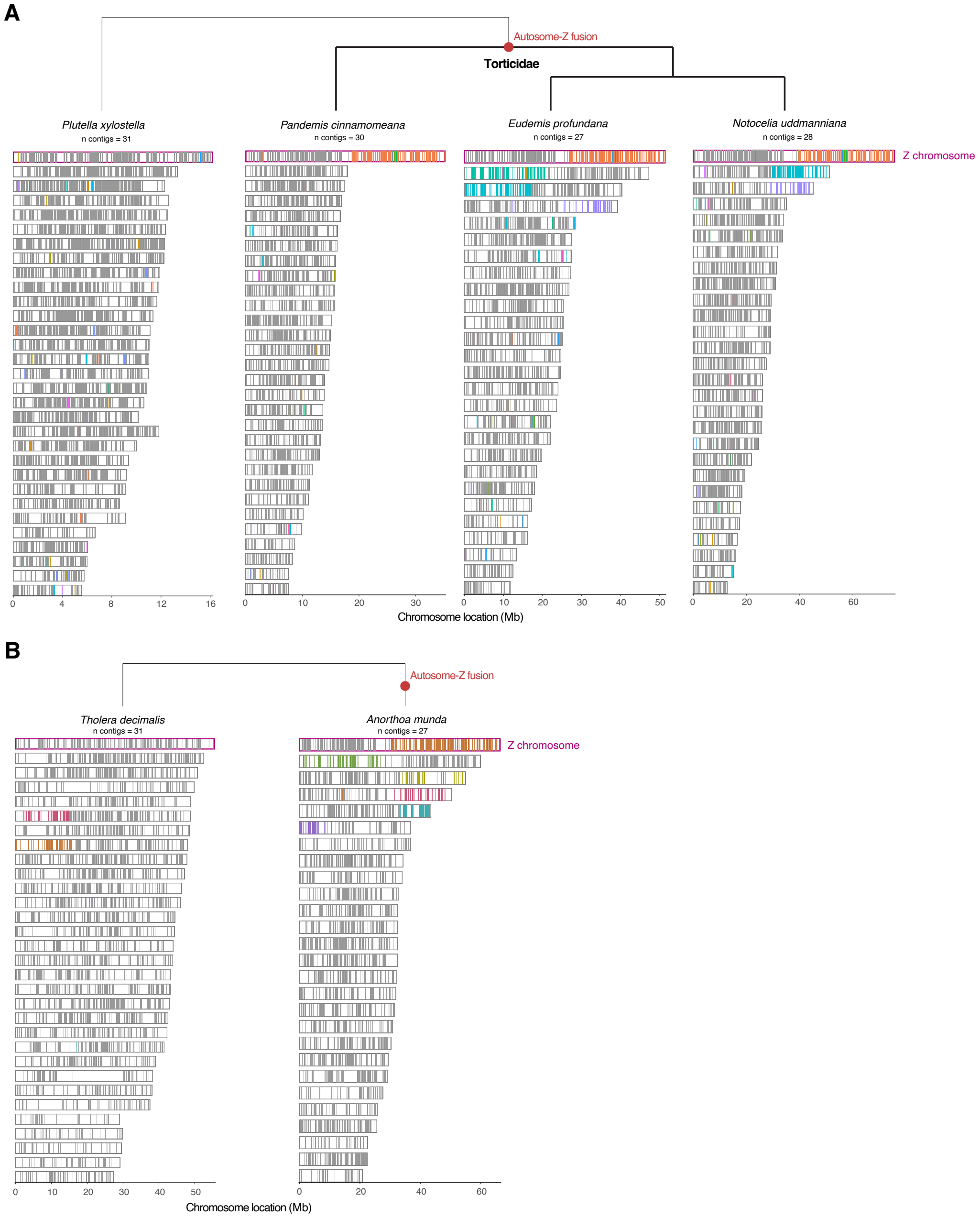


**Supplemental Figure S5: Autosome-Z chromosome fusion events in Tortricidae and *Anorthoa munda*.** Orthologous BUSCO genes between outgroup and ingroup species are painted onto each chromosome using lepbuscopainter (`https://github.com/charlottewright/lep_busco_painter`). For each species, each chromosome is shown as a rectangle. A grey bar is given for shared orthologous BUSCO genes in the same position as the outgroup species, and a coloured bar is shown for BUSCO genes in an alternative chromosome location compared to the outgroup. This is shown for **(A)** three representative species in Tortricidae compared to *Plutella xylostella* as the outgroup, and **(B)** *Anorthoa munda* compared to *Tholera decimalis* as the outgroup species. In both **A** and **B**, the top chromosome rectangle represents the Z chromosome in all species (highlighted with a pink box) and shows an autosome-Z chromosome fusion in the ingroup species.

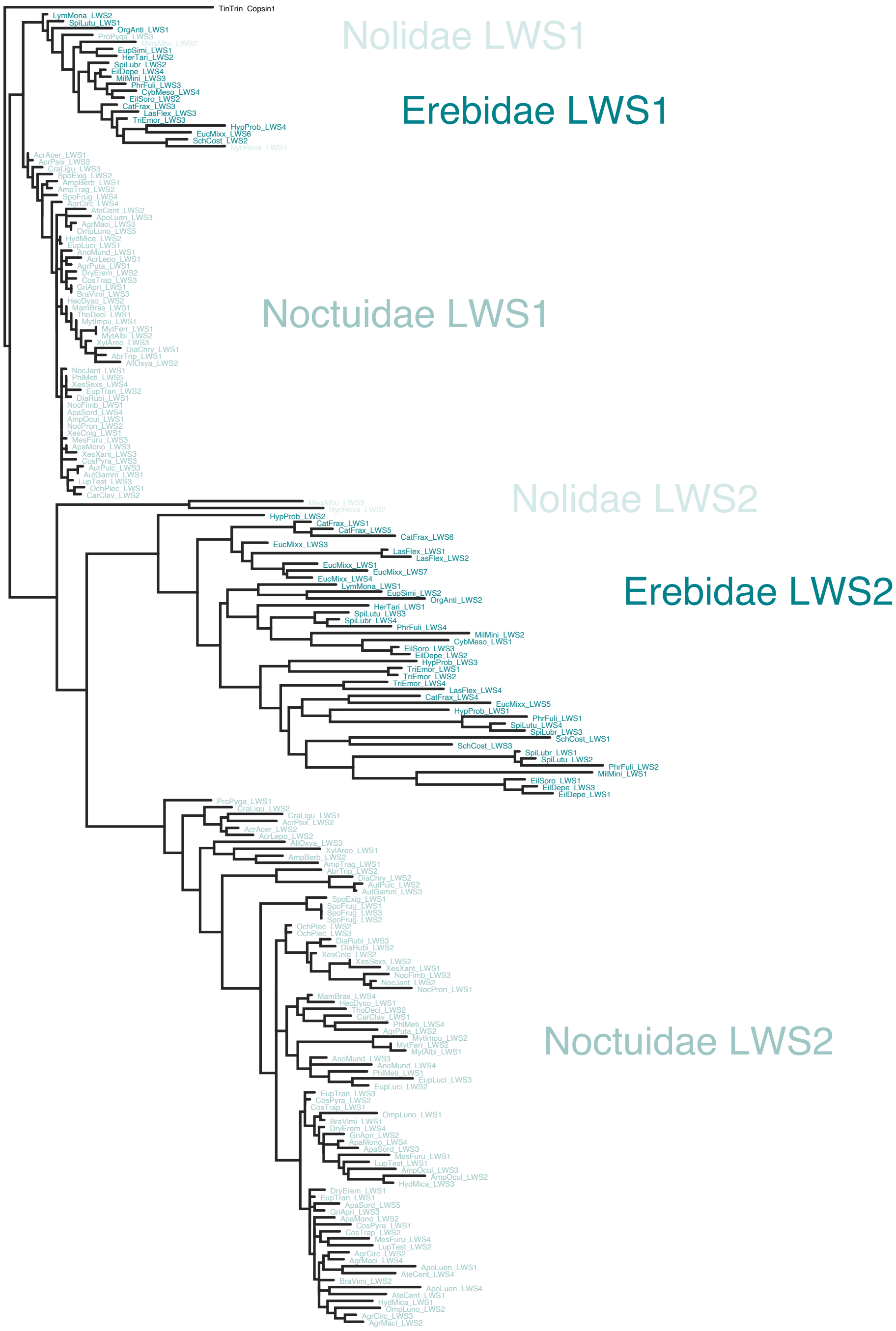


**Supplemental Figure S6: Noctuoidea LW opsin gene tree.** Gene tree of all LW opsin copies present in species within the superfamily Noctuoidea (Noldiae, Erebidae, and Noctuidae). Each family has a unique colour, and each clade corresponding to an LW gene is labelled. LWS1 represents the parent LW copy in these species, while the LW2 clades represent the ancestrally duplicated LW copy.

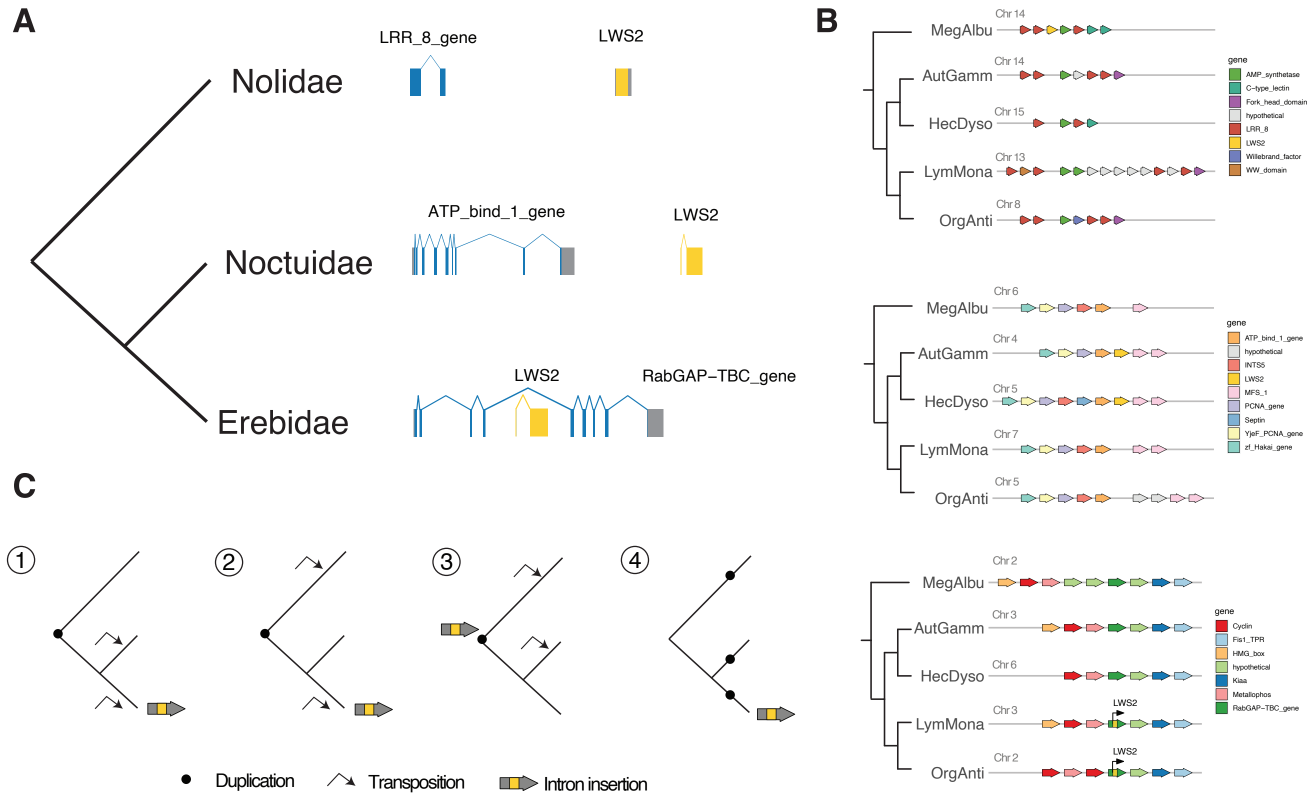


**Supplemental Figure S7: Summary of LWS2 paralog genomic locations and duplication history.** **(A)** Topology of each family within the Noctuoidea superfamily (left) and the corresponding representative genomic location of the LWS2 copy in each family (right). The LWS2 copy is shown in yellow, while the genes located within the syntenic region in each family is shown in blue and labelled according to the functional domain contained within that gene. **(B)** Family specific syntenic clusters surrounding the LWS2 gene. For each family specific LWS2 synetic block (top, middle, and bottom), the surrounding genes are shown for *Meganola albula* (Nolidae), *Autographa gamma* (Noctuidae), *Hecatera dysodea* (Noctuidae), *Lymantria monacha* (Erebidae), and *Orgyia antiqua* (Erebidea). The syntenic genes surrounding the LWS2 copy are coloured and the corresponding colour and gene name are given in the legend. **(C)** The hypothetical modes of LWS2 origination and subsequent genome translocations are provides along the topology which corresponds to that in (A).

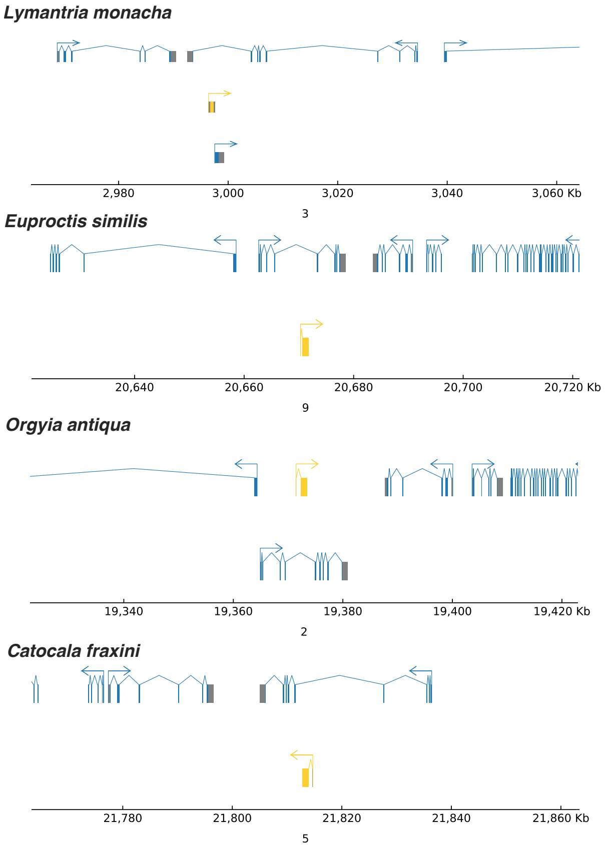


**Supplemental Figure S8: Genomic locations of LWS2 paralog in Erebidae species.** The location of the LWS2 gene within the intron of another gene, as well as the surrounding syntenic genes, is shown for 8 species from the Erebidae family. In each case the LWS2 gene is shown in yellow, and the surrounding genes are shown in blue. The chromosome and position within the chromosome is given for each one.

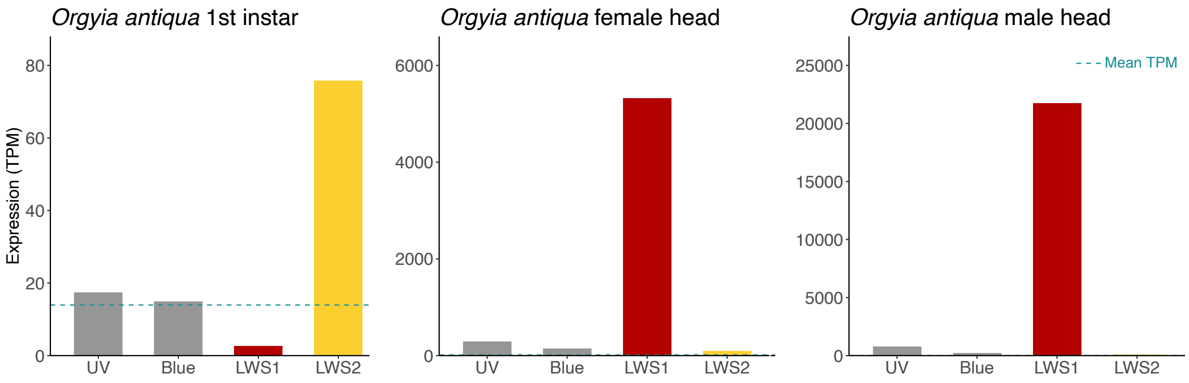


**Supplemental Figure S9: TPM values for opsin genes in different lifestages of Vapourer moth (*Orgyia antiqua*).** TPM values are given for each of the visual opsin genes (UV, Blue, and LW) in *Orgyia antiqua* first instar larvae (left), female head (middle), and male head (right). In each case, the bar corresponding to the parent LWS1 gene is coloured red and the bar corresponding to the duplicated LWS2 gene is coloured yellow. The average TPM values for each sample is shown by a broken blue line.

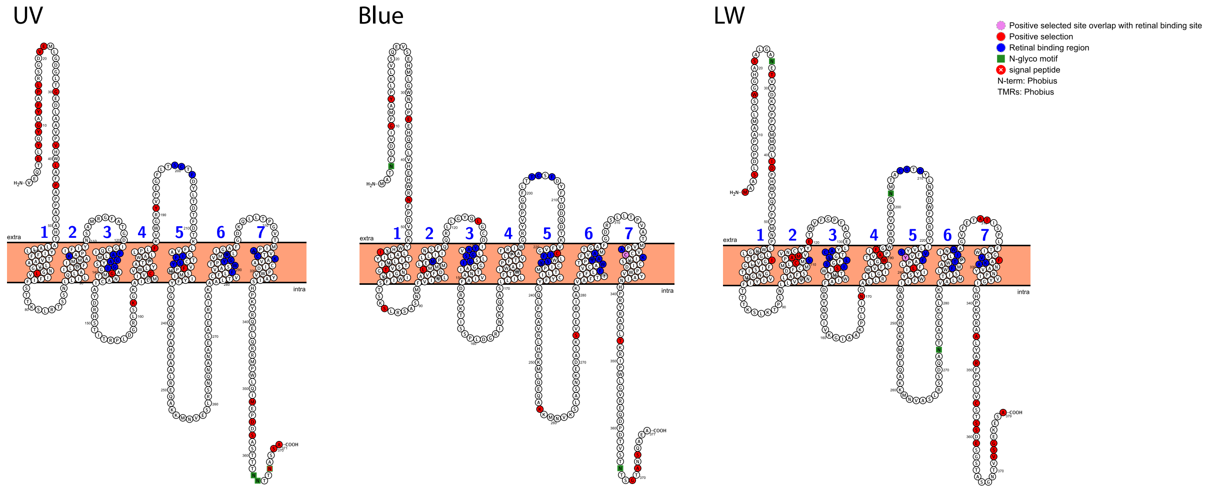


**Supplemental Figure S10: Opsin structure with retinal binding sites and overlap with positively selected sites.** PHOBIUS transmembrane predictions for each of the three visual opsin genes (UV, Blue, LW) used in the selective pressure analysis. Amino acid sites within 5Å of the chromophore binding pocket (coloured blue) were identified by homology modelling against the crystal structure of Jumping Spider Rhodopsin-1. Sites found to be under positive selection (branch site test) in any of the transition branches from noctural to diurnal species are highlighted by the red colour. Sites where positive selection was found to overlap with retinal binding sites are coloured purple.

### Supplementary Tables

**Table S1: Species sampled and genome source information.**

| **Species name** | **Short name** | **GenBank accession** | **Project ID** |
|---|---|---|---|
| Micropterix aruncella | MicArun | GCA_944548615 | PRJEB40665 |
| Neomicropteryx cornuta | NeoCorn | GCA_020383195 | PRJNA731916 |
| Incurvaria masculella | IncMasc | GCA_946894095 | PRJEB40665 |
| Nematopogon swammerdamellus | NemSwam | GCA_946902875 | PRJEB40665 |
| Tinea semifulvella | TinSemi | GCA_910589645 | PRJEB40665 |
| Tinea trinotella | TinTrin | GCA_905220615 | PRJEB40665 |
| Yponomeuta sedellus | YpoSede | GCA_934045075 | PRJEB40665 |
| Plutella xylostella | PluXylo | GCA_932276165 | PRJEB40665 |
| Ypsolopha scabrella | YpsScab | GCA_910592155 | PRJEB40665 |
| Ypsolopha sequella | YpsSequ | GCA_934047225 | PRJEB40665 |
| Pandemis cinnamomeana | PanCinn | GCA_932294345 | PRJEB40665 |
| Acleris emargana | AclEmar | GCA_927399475 | PRJEB40665 |
| Acleris literana | AclLite | GCA_946894065 | PRJEB40665 |
| Acleris sparsana | AclSpar | GCA_923062465 | PRJEB40665 |
| Eudemis profundana | EudProf | GCA_947034925 | PRJEB40665 |
| Hedya salicella | HedSali | GCA_905404275 | PRJEB40665 |
| Apotomis betuletana | ApoBetu | GCA_932273695 | PRJEB40665 |
| Apotomis turbidana | ApoTurb | GCA_905147355 | PRJEB40665 |
| Cydia splendana | CydSple | GCA_910591565 | PRJEB40665 |
| Pammene aurita | PamAuri | GCA_947086415 | PRJEB40665 |
| Pammene fasciana | PamFasc | GCA_911728535 | PRJEB40665 |
| Notocelia uddmanniana | NotUddm | GCA_905163555 | PRJEB40665 |
| Epinotia nisella | EpiNise | GCA_932294315 | PRJEB40665 |
| Epinotia bilunana | EpiBilu | GCA_947049275 | PRJEB40665 |
| Epinotia demarniana | EpiDema | GCA_945867215 | PRJEB40665 |
| Apoda limacodes | ApoLima | GCA_946406115 | PRJEB40665 |
| Zygaena filipendulae | ZygFili | GCA_907165275 | PRJEB40665 |
| Zeuzera pyrina | ZeuPyri | GCA_907165235 | PRJEB40665 |
| Sesia apiformis | SesApif | GCA_914767545 | PRJEB40665 |
| Sesia bembeciformis | SesBemb | GCA_943735995 | PRJEB40665 |
| Bembecia ichneumoniformis | BemIchn | GCA_910589475 | PRJEB40665 |
| Synanthedon andrenaeformis | SynAndr | GCA_936446665 | PRJEB40665 |
| Synanthedon myopaeformis | SynMyop | GCA_944738685 | PRJEB40665 |
| Synanthedon formicaeformis | SynForm | GCA_945859745 | PRJEB40665 |
| Synanthedon vespiformis | SynVesp | GCA_918317495 | PRJEB40665 |
| Emmelina monodactyla | EmmMono | GCA_916618145 | PRJEB40665 |
| Marasmarcha lunaedactyla | MarLuna | GCA_923062675 | PRJEB40665 |
| Stenoptilia bipunctidactyla | SteBipu | GCA_944452665 | PRJEB40665 |
| Papilio machaon | PapMach | GCA_912999745 | PRJEB40665 |
| Erynnis tages | EryTage | GCA_905147235 | PRJEB40665 |
| Pyrgus malvae | PyrMalv | GCA_911387765 | PRJEB40665 |

| | | | |
|---|---|---|---|
| Carterocephalus palaemon | CarPala | GCA_944567765 | PRJEB40665 |
| Thymelicus sylvestris | ThySylv | GCA_911387775 | PRJEB40665 |
| Hesperia comma | HesComm | GCA_905404135 | PRJEB40665 |
| Ochlodes sylvanus | OchSylv | GCA_905404295 | PRJEB40665 |
| Leptidea sinapis | LepSina | GCA_905404315 | PRJEB40665 |
| Colias croceus | ColCroc | GCA_905220415 | PRJEB40665 |
| Anthocharis cardamines | AntCard | GCA_905404175 | PRJEB40665 |
| Aporia crataegi | ApoCrat | GCA_912999735 | PRJEB40665 |
| Pieris rapae | PieRapa | GCA_905147795 | PRJEB40665 |
| Pieris brassicae | PieBras | GCA_905147105 | PRJEB40665 |
| Pieris napi | PieNapi | GCA_905231885 | PRJEB40665 |
| Lycaena phlaeas | LycPhla | GCA_905333005 | PRJEB40665 |
| Celastrina argiolus | CelArgi | GCA_905187575 | PRJEB40665 |
| Glaucopsyche alexis | GlaAlex | GCA_905404095 | PRJEB40665 |
| Plebejus argus | PleArgu | GCA_905404155 | PRJEB40665 |
| Cyaniris semiargus | CyaSemi | GCA_905187585 | PRJEB40665 |
| Aricia agestis | AriAges | GCA_944452695 | PRJEB40665 |
| Aricia artaxerxes | AriArta | GCA_937612035 | PRJEB40665 |
| Polyommatus icarus | PolIcar | GCA_937595015 | PRJEB40665 |
| Lysandra bellargus | LysBell | GCA_905333045 | PRJEB40665 |
| Lysandra coridon | LysCori | GCA_905220515 | PRJEB40665 |
| Bicyclus anynana | BicAnyn | GCA_947172395 | PRJEB40665 |
| Lasiommata megera | LasMege | GCA_928268935 | PRJEB40665 |
| Pararge aegeria | ParAege | GCA_905163445 | PRJEB40665 |
| Hipparchia semele | HipSeme | GCA_933228805 | PRJEB40665 |
| Melanargia galathea | MelGala | GCA_920104075 | PRJEB40665 |
| Aphantopus hyperantus | AphHype | GCA_902806685 | PRJEB40665 |
| Maniola jurtina | ManJurt | GCA_905333055 | PRJEB40665 |
| Erebia aethiops | EreAeth | GCA_923060345 | PRJEB40665 |
| Erebia ligea | EreLige | GCA_917051295 | PRJEB40665 |
| Limenitis camilla | LimCami | GCA_905147385 | PRJEB40665 |
| Boloria selene | BolSele | GCA_905231865 | PRJEB40665 |
| Fabriciana adippe | FabAdip | GCA_905404265 | PRJEB40665 |
| Mellicta athalia | MelAtha | GCA_905220545 | PRJEB40665 |
| Melitaea cinxia | MelCinx | GCA_905220565 | PRJEB40665 |
| Vanessa atalanta | VanAtal | GCA_905147765 | PRJEB40665 |
| Vanessa cardui | VanCard | GCA_905220365 | PRJEB40665 |
| Nymphalis polychloros | NymPoly | GCA_905220585 | PRJEB40665 |
| Inachis io | AglIo | GCA_905147045 | PRJEB40665 |
| Nymphalis urticae | AglUrti | GCA_905147175 | PRJEB40665 |
| Agonopterix subpropinquella | AgoSubp | GCA_922987775 | PRJEB40665 |
| Carcina quercana | CarQuer | GCA_910589575 | PRJEB40665 |
| Hyposmocoma kahamanoa | HypKaha | GCA_003589595 | PRJEB40665 |
| Coleophora flavipennella | ColFlav | GCA_947284805 | PRJEB40665 |
| Esperia sulphurella | EspSulp | GCA_947086405 | PRJEB40665 |
| Blastobasis adustella | BlaAdus | GCA_907269095 | PRJEB40665 |
| Blastobasis lacticolella | BlaLact | GCA_905147135 | PRJEB40665 |

| | | | |
|---|---|---|---|
| Acrobasis suavella | AcrSuav | GCA_943193695 | PRJEB40665 |
| Apomyelois bistriatella | ApoBist | GCA_947044815 | PRJEB40665 |
| Endotricha flammealis | EndFlam | GCA_905163395 | PRJEB40665 |
| Hypsopygia costalis | HypCost | GCA_937001555 | PRJEB40665 |
| Acentria ephemerella | AceEphe | GCA_943193645 | PRJEB40665 |
| Parapoynx stratiotata | ParStra | GCA_910589355 | PRJEB40665 |
| Calamotropha paludella | CalPalu | GCA_927399485 | PRJEB40665 |
| Chrysoteuchia culmella | ChrCulm | GCA_910589605 | PRJEB40665 |
| Agriphila geniculea | AgrGeni | GCA_943789525 | PRJEB40665 |
| Agriphila tristella | AgrTris | GCA_928269145 | PRJEB40665 |
| Drepana falcataria | DreFalc | GCA_945859725 | PRJEB40665 |
| Watsonalla binaria | WatBina | GCA_929442735 | PRJEB40665 |
| Habrosyne pyritoides | HabPyri | GCA_907165245 | PRJEB40665 |
| Thyatira batis | ThyBati | GCA_905147785 | PRJEB40665 |
| Saturnia pavonia | SatPavo | GCA_947532125 | PRJEB40665 |
| Deilephila porcellus | DeiPorc | GCA_905220455 | PRJEB40665 |
| Hemaris fuciformis | HemFuci | GCA_907164795 | PRJEB40665 |
| Laothoe populi | LaoPopu | GCA_905220505 | PRJEB40665 |
| Mimas tiliae | MimTili | GCA_905332985 | PRJEB40665 |
| Ligdia adustata | LigAdus | GCA_947049295 | PRJEB40665 |
| Macaria notata | MacNota | GCA_927399415 | PRJEB40665 |
| Biston betularia | BisBetu | GCA_905404145 | PRJEB40665 |
| Erannis defoliaria | EraDefo | GCA_905404285 | PRJEB40665 |
| Peribatodes rhomboidaria | PerRhom | GCA_911728515 | PRJEB40665 |
| Agriopis aurantiaria | AgrAura | GCA_914767915 | PRJEB40665 |
| Agriopis marginaria | AgrMarg | GCA_932305915 | PRJEB40665 |
| Apeira syringaria | ApeSyri | GCA_934044485 | PRJEB40665 |
| Selenia dentaria | SelDent | GCA_917880725 | PRJEB40665 |
| Alsophila aescularia | AlsAesc | GCA_946251855 | PRJEB40665 |
| Campaea margaritaria | CamMarg | GCA_912999815 | PRJEB40665 |
| Hylaea fasciaria | HylFasc | GCA_905147375 | PRJEB40665 |
| Opisthograptis luteolata | OpiLute | GCA_931315375 | PRJEB40665 |
| Crocallis elinguaria | CroElin | GCA_907269065 | PRJEB40665 |
| Ennomos fuscantarius | EnnFusc | GCA_905220475 | PRJEB40665 |
| Ennomos quercinarius | EnnQuer | GCA_910589525 | PRJEB40665 |
| Idaea aversata | IdaAver | GCA_907269075 | PRJEB40665 |
| Aplocera efformata | AplEffo | GCA_921293045 | PRJEB40665 |
| Lobophora halterata | LobHalt | GCA_932526365 | PRJEB40665 |
| Gymnoscelis rufifasciata | GymRufi | GCA_929108375 | PRJEB40665 |
| Eupithecia centaureata | EupCent | GCA_944547425 | PRJEB40665 |
| Eupithecia abbreviata | EupAbbr | GCA_943735965 | PRJEB40665 |
| Eupithecia dodoneata | EupDodo | GCA_947044415 | PRJEB40665 |
| Eupithecia exiguata | EupExig | GCA_947086465 | PRJEB40665 |
| Eupithecia vulgata | EupVulg | GCA_946478455 | PRJEB40665 |
| Operophtera brumata | OpeBrum | GCA_932527175 | PRJEB40665 |
| Philereme vetulata | PhiVetu | GCA_918857605 | PRJEB40665 |
| Hydriomena furcata | HydFurc | GCA_912999785 | PRJEB40665 |

| | | | |
|---|---|---|---|
| Xanthorhoe spadicearia | XanSpad | GCA_947086425 | PRJEB40665 |
| Ecliptopera silaceata | EclSila | GCA_932527185 | PRJEB40665 |
| Eulithis prunata | EulPrun | GCA_918843925 | PRJEB40665 |
| Electrophaes corylata | EleCory | GCA_947095575 | PRJEB40665 |
| Chloroclysta siterata | ChlSite | GCA_932294275 | PRJEB40665 |
| Thera britannica | TheBrit | GCA_939531255 | PRJEB40665 |
| Clostera curtula | CloCurt | GCA_905475355 | PRJEB40665 |
| Furcula furcula | FurFurc | GCA_911728495 | PRJEB40665 |
| Phalera bucephala | PhaBuce | GCA_905147815 | PRJEB40665 |
| Pheosia gnoma | PheGnom | GCA_905404115 | PRJEB40665 |
| Pheosia tremula | PheTrem | GCA_905333125 | PRJEB40665 |
| Ptilodon capucinus | PtiCapu | GCA_914767695 | PRJEB40665 |
| Notodonta dromedarius | NotDrom | GCA_905147325 | PRJEB40665 |
| Notodonta ziczac | NotZicz | GCA_918843915 | PRJEB40665 |
| Meganola albula | MegAlbu | GCA_936450015 | PRJEB40665 |
| Nycteola revayana | NycReva | GCA_947037095 | PRJEB40665 |
| Lymantria monacha | LymMona | GCA_905163515 | PRJEB40665 |
| Euproctis similis | EupSimi | GCA_905147225 | PRJEB40665 |
| Orgyia antiqua | OrgAnti | GCA_916999025 | PRJEB40665 |
| Catocala fraxini | CatFrax | GCA_930367265 | PRJEB40665 |
| Euclidia mi | EucMi | GCA_944739405 | PRJEB40665 |
| Schrankia costaestrigalis | SchCost | GCA_905475405 | PRJEB40665 |
| Laspeyria flexula | LasFlex | GCA_905147015 | PRJEB40665 |
| Trisateles emortualis | TriEmor | GCA_947095525 | PRJEB40665 |
| Hypena proboscidalis | HypProb | GCA_905147285 | PRJEB40665 |
| Herminia tarsipennalis | HerTars | GCA_945859575 | PRJEB40665 |
| Phragmatobia fuliginosa | PhrFuli | GCA_932526445 | PRJEB40665 |
| Spilosoma lubricipeda | SpiLubr | GCA_905220595 | PRJEB40665 |
| Spilarctia lutea | SpiLute | GCA_916048165 | PRJEB40665 |
| Miltochrista miniata | MilMini | GCA_933228765 | PRJEB40665 |
| Cybosia mesomella | CybMeso | GCA_946251805 | PRJEB40665 |
| Eilema depressum | EilDepr | GCA_914767945 | PRJEB40665 |
| Eilema sororculum | EilSoro | GCA_914829495 | PRJEB40665 |
| Abrostola tripartita | AbrTrip | GCA_946251915 | PRJEB40665 |
| Diachrysia chrysitis | DiaChry | GCA_932294365 | PRJEB40665 |
| Autographa gamma | AutGamm | GCA_905146925 | PRJEB40665 |
| Autographa pulchrina | AutPulc | GCA_905475315 | PRJEB40665 |
| Protodeltote pygarga | ProPyga | GCA_936450705 | PRJEB40665 |
| Allophyes oxyacanthae | AllOxya | GCA_932294325 | PRJEB40665 |
| Craniophora ligustri | CraLigu | GCA_905163465 | PRJEB40665 |
| Acronicta psi | AcrPsi | GCA_946251955 | PRJEB40665 |
| Acronicta aceris | AcrAcer | GCA_910591435 | PRJEB40665 |
| Acronicta leporina | AcrLepo | GCA_947256265 | PRJEB40665 |
| Xylocampa areola | XylAreo | GCA_935421205 | PRJEB40665 |
| Amphipyra berbera | AmpBerb | GCA_910594945 | PRJEB40665 |
| Amphipyra tragopoginis | AmpTrag | GCA_905220435 | PRJEB40665 |
| Spodoptera exigua | SpoExig | GCA_902829305 | PRJEB36598 |

| | | | |
|---|---|---|---|
| Spodoptera frugiperda | SpoFrug | GCA_011064685 | PRJNA590312 |
| Caradrina clavipalpis | CarClav | GCA_932526535 | PRJEB40665 |
| Anorthoa munda | AnoMund | GCA_945859665 | PRJEB40665 |
| Tholera decimalis | ThoDeci | GCA_943138885 | PRJEB40665 |
| Hecatera dysodea | HecDyso | GCA_905332915 | PRJEB40665 |
| Mamestra brassicae | MamBras | GCA_905163435 | PRJEB40665 |
| Mythimna impura | MytImpu | GCA_905147345 | PRJEB40665 |
| Mythimna albipuncta | MytAlbi | GCA_929112965 | PRJEB40665 |
| Mythimna ferrago | MytFerr | GCA_910589285 | PRJEB40665 |
| Agrotis puta | AgrPuta | GCA_943136025 | PRJEB40665 |
| Ochropleura plecta | OchPlec | GCA_905475445 | PRJEB40665 |
| Diarsia rubi | DiaRubi | GCA_932274075 | PRJEB40665 |
| Noctua pronuba | NocPron | GCA_905220335 | PRJEB40665 |
| Noctua fimbriata | NocFimb | GCA_905163415 | PRJEB40665 |
| Noctua janthe | NocJant | GCA_910589295 | PRJEB40665 |
| Xestia c-nigrum | XesCnig | GCA_916618015 | PRJEB40665 |
| Xestia sexstrigata | XesSexs | GCA_941918905 | PRJEB40665 |
| Xestia xanthographa | XesXant | GCA_905147715 | PRJEB40665 |
| Euplexia lucipara | EupLuci | GCA_921972225 | PRJEB40665 |
| Phlogophora meticulosa | PhlMeti | GCA_905147745 | PRJEB40665 |
| Atethmia centrago | AteCent | GCA_905333075 | PRJEB40665 |
| Cosmia pyralina | CosPyra | GCA_946251885 | PRJEB40665 |
| Cosmia trapezina | CosTrap | GCA_905163495 | PRJEB40665 |
| Eupsilia transversa | EupTran | GCA_914767815 | PRJEB40665 |
| Omphaloscelis lunosa | OmpLuno | GCA_916610215 | PRJEB40665 |
| Agrochola circellaris | AgrCirc | GCA_914767755 | PRJEB40665 |
| Agrochola macilenta | AgrMaci | GCA_916701695 | PRJEB40665 |
| Brachylomia viminalis | BraVimi | GCA_937001585 | PRJEB40665 |
| Dryobotodes eremita | DryErem | GCA_917490735 | PRJEB40665 |
| Griposia aprilina | GriApri | GCA_916610205 | PRJEB40665 |
| Aporophyla lueneburgensis | ApoLuen | GCA_932294355 | PRJEB40665 |
| Apamea monoglypha | ApaMono | GCA_911387795 | PRJEB40665 |
| Apamea sordens | ApaSord | GCA_945859715 | PRJEB40665 |
| Amphipoea oculea | AmpOcul | GCA_945859645 | PRJEB40665 |
| Hydraecia micacea | HydMica | GCA_914767645 | PRJEB40665 |
| Luperina testacea | LupTest | GCA_927399505 | PRJEB40665 |
| Mesoligia furuncula | MesFuru | GCA_916614155 | PRJEB40665 |
